# Benchmarking imputation methods for network inference using a novel method of synthetic scRNA-seq data generation

**DOI:** 10.1101/2021.10.13.464275

**Authors:** Ayoub Lasri, Vahid Shahrezaei, Marc Sturrock

## Abstract

Single cell RNA-sequencing (scRNA-seq) has very rapidly become the new workhorse of modern biology providing an unprecedented global view on cellular diversity and heterogeneity. In particular, the structure of gene-gene expression correlation contains information on the underlying gene regulatory networks. However, interpretation of scRNA-seq data is challenging due to specific experimental error and biases that are unique to this kind of data including drop-out (or technical zeros). To deal with this problem several methods for imputation of zeros for scRNA-seq have been developed. However, it is not clear how these processing steps affect inference of genetic networks from single cell data. Here, we introduce Biomodelling.jl, a tool for generation of synthetic scRNA-seq data using multiscale modelling of stochastic gene regulatory networks in growing and dividing cells. Our tool produces realistic transcription data with a known ground truth network topology that can be used to benchmark different approaches for gene regulatory network inference. Using this tool we investigate the impact of different imputation methods on the performance of several network inference algorithms. Biomodelling.jl provides a versatile and useful tool for future development and benchmarking of network inference approaches using scRNA-seq data.

## 1 Introduction

A gene regulatory network (GRN) or genetic network (GN) refers to a collection of interacting genes in a cell which regulate each other indirectly through interaction of their protein expression products and regulatory parts of DNA and with other signalling systems in the cell, thereby governing the rates at which genes in the cell are transcribed into mRNA [1]. GRNs can be represented as graphs or networks, where the nodes of the network are genes and the edges between nodes represent gene interactions through which the products of one gene affect those of another. These interactions can be activating, with an increase in the expression of one leading to an increase in the other, or inhibiting, with an increase in one leading to a decrease in the other. Learning the structure and behaviour of GRNs is a fundamental problem in biology since many cellular processes, such as the cell cycle, cellular differentiation, and apoptosis are tightly controlled by GRNs. Hence the elucidation of these GRNs is of critical importance in many fields such as medicine and systems biology, however progress in deciphering them has been slow.

In recent years, high-throughput sequencing methods have revolutionised the entire field of biology. The opportunity to study entire transcriptomes in great detail using RNA sequencing (RNA-seq) has catalysed many important discoveries and is now a routine method in biomedical research. However, RNA-seq is typically performed in “bulk”, and the data represent an average of gene expression patterns across thousands to millions of cells. This averaging obscures biologically relevant differences between cells and limits the possible downstream analyses. Single-cell RNA-seq (scRNA-seq) represents an approach to overcome this problem [2]. By isolating single cells, capturing their transcripts, and generating sequencing libraries in which the transcripts are mapped to individual cells, scRNA-seq allows assessment of fundamental biological properties of cell populations and biological systems at unprecedented resolution.

Unlike traditional profiling methods that assess bulk populations, scRNA-seq offers an insight into biologically relevant cell-to-cell variations in gene expression. This includes understanding the tumour microenvironment [3] by revealing complex and rare populations [4], facilitating the tracking of trajectories of cell lineages [5] and providing insights into heterogeneity of stress response in microbes [6]. As we will explore in this paper, it can facilitate the inference of GRNs [7]. Nevertheless, many factors contribute to the rise of analysis challenges when dealing with scRNA-seq data, such factors can be divided into two main classes: technical variation (e.g. batch effect, cell specific capture efficiency, amplification bias and dropout events) and biological variation (e.g. stochastic gene expression, cell differentiation, environmental niche and cell cycle).

Over the last decade many inference methods have been developed to harness the available high-throughput data such as the RNA-seq data to uncover regulatory interactions in GRNs. GRN inference is usually performed on measurements of gene-gene correlation, mutual information or regression models that can be obtained from bulk RNA-seq data across multiple conditions or perturbations or scRNA-seq across many cells. If a co-expression between two genes is detected, while considering the expression of all others genes (conditional information), these genes are said to have a regulatory relationship. Several methods have been developed specifically for scRNA-seq [8, 9] but some reviews and benchmarking studies have shown that both bulk and single cell methods perform poorly on scRNA-seq data [10, 11]. For more accurate GRN reconstruction several authors have remarked that preprocessing the data is important, mostly due to the sparse nature of the data [12, 13]. Among different preprocessing steps, normalisation and imputation is of particular importance. In order to distinguish between biological and technical zeros (drop-out events), several imputation methods have been developed [14, 15, 16, 17, 18, 19] and compared in benchmark studies [20, 21]. The imputation step is often integrated with normalisation and other downstream analysis as implemented in these methods [22, 19]. However, how imputation affects gene-gene correlations is not entirely clear although there have been some studies that have suggested that performing imputation improves the estimation of gene-gene correlations [18, 23]. So, there seems to be some potential for using imputation methods to improve GRN inference from scRNA-seq data.

While many methods have been developed for inference of gene regulatory networks, evaluating the performance of these methods remains challenging due to lack of appropriate benchmarks. In general, there are three main strategies to generate benchmark networks. A first strategy consists in evaluating network predictions made by reverse engineering algorithms on well-studied *in vivo* pathways from model organisms [24, 25]. However, those networks are incomplete maps of the physical interactions in the cell that are responsible for cellular functions and using them as benchmarks will inevitably lead to errors when evaluating network predictions. Another strategy consists of genetically engineering synthetic in vivo networks [26, 27]. The main drawback of this strategy is that only a few small networks are available. The third strategy consists of developing in *silico* gene regulatory networks that can be simulated to produce synthetic gene expression data that can be used in bench-marking. The simulation of *in silico* networks has the advantages of being fast, easily reproducible and less expensive than biological experiments and the ground truth is exactly known. However, for the synthetic data to be useful, it should have a realistic assumptions and statistical properties for the underlying GRN topology and gene expression.

Benchmark synthetic data generators such as “artificial gene networks” [28] aim to produce *in silico* gene networks exhibiting topological properties observed in biological networks using Erdös-Renyi, Watts-Strogatz (small-world) or Albert-Barabási (scale-free) random graph models. Other approaches have been taken in SynTReN [29] and [30] where general network structures were created by extracting parts of known *in vivo* regulatory network structures. These approaches have the advantage of capturing several structural properties observed in in vivo network structures. In order to produce temporal gene expression data, the generated structures are often made using dynamical models of gene regulation. Systems of non-linear ordinary differential equations (ODEs) are widely used [31]. As current high-throughput technologies that simultaneously monitor protein expression are limited, some benchmark generators consider mRNA as a proxy for protein expression and thus do not model translation independently of transcription [30, 29]. Protein expression in general does not correlate well with mRNA expression in many biological systems [32]. To overcome this, several benchmark synthetic data generators have accounted for transcription and translation explicitly such as RENCO [33], GeNGe [34] and GREN-DEL [35]. GeneNetWeaver has become a commonly used tool in recent years to generate gene expression data and GRN model evaluations [36]. For instance, it was selected to generate the “gold standard” networks for the DREAM4 and DREAM5 network inference challenges, as well as other publications that conducted comparisons of network modelling approaches [37, 38, 39]. GeneNetWeaver uses chemical langevin equations to simulate stochastic gene expression and allows for both independent (‘additive’) and synergistic (‘multiplicative’) interactions. Among methods that creates statistically realistic synthetic scRNA-seq data generation method is splatter [40]. Splatter implements six different simulation models ranging from a simple negative Binomial model to a more sophisticated gamma-Poisson hierarchical model, however, it assumes no correlation in expression among different genes. Finally, MeSCoT was released recently which is a synthetic data generator developed in MATLAB for the detailed simulation of genes’ regulatory interactions for variable genomic architectures which can also produce a complete set of transcriptional and translational data together with simulated quantitative trait values [41]. So, while there are several *in silico* methods available for simulating gene expression data, currently no method produces synthetic scRNA-seq data with realistic expression statistics as expected by stochastic gene expression and scRNA-seq protocols.

In this paper, we propose a novel in *silico* tool written purely in Julia [42] to generate synthetic scRNA-seq data suitable for benchmarking GRN inference methods, Biomodelling.jl*^1^. Our method uses an agent-based method to couple stochastic simulations of realistic GRNs in a population of growing and dividing cells. We couple cell size to transcription as has recently been observed in different cellular systems [43] and include translation, binomial partitioning of molecules upon cell division and capture efficiency of the scRNA-seq steps. Here, we used Biomodelling.jl to systematically benchmark the impact of different imputation methods on the performance of network inference algorithms.

The format of this paper is as follows. We begin in section 2 by introducing our method of synthetic data generation as well as the different imputation methods and network inference methods we wish to assess. We then begin section 3 by presenting a toy 5 gene example as an exemplar of our method and use it to illustrate the central problem of overcoming the negative impact of downsampling on network inference. Next we show that the network inference methods perform better on sparser data before going onto show how the different imputation methods and network inference methods perform using realistic scale-free topologies. We show that multiplicative regulation is the most challenging for accurate network inference. We then show that the best choice of imputation method for accurate inference depends on the choice of inference method. Finally we show that the number of combination reactions (where a gene has multiple regulators) considered rather than the size of the network determines overall performance. We end with a discussion in section 4 and make some recommendations for how best to pre-process scRNA-seq data for network inference.

## 2 Methods

### 2.1 Biomodelling.jl

*Biomodelling.jl* is a tool for multiscale agent-based modelling of scRNA-seq data that simulates stochastic gene expression in a population of single cells that are growing and dividing, written in the Julia programming language. The unique feature of *Biomodelling.jl* is that it can generate synthetic scRNA-seq from a known underlying gene regulatory network including global transcription-cell volume relationships. In Figure 1, we describe the main steps in order to generate synthetic ground truth (GT) data using *Biomodelling.jl*, which is available to the community as open source software. The gene-gene correlation that is exhibited in the Biomodelling.jl synthetic data provides benchmarking data for testing the efficiency of network inference methods. Details about each step are given in the following sections.

**Figure 1:**
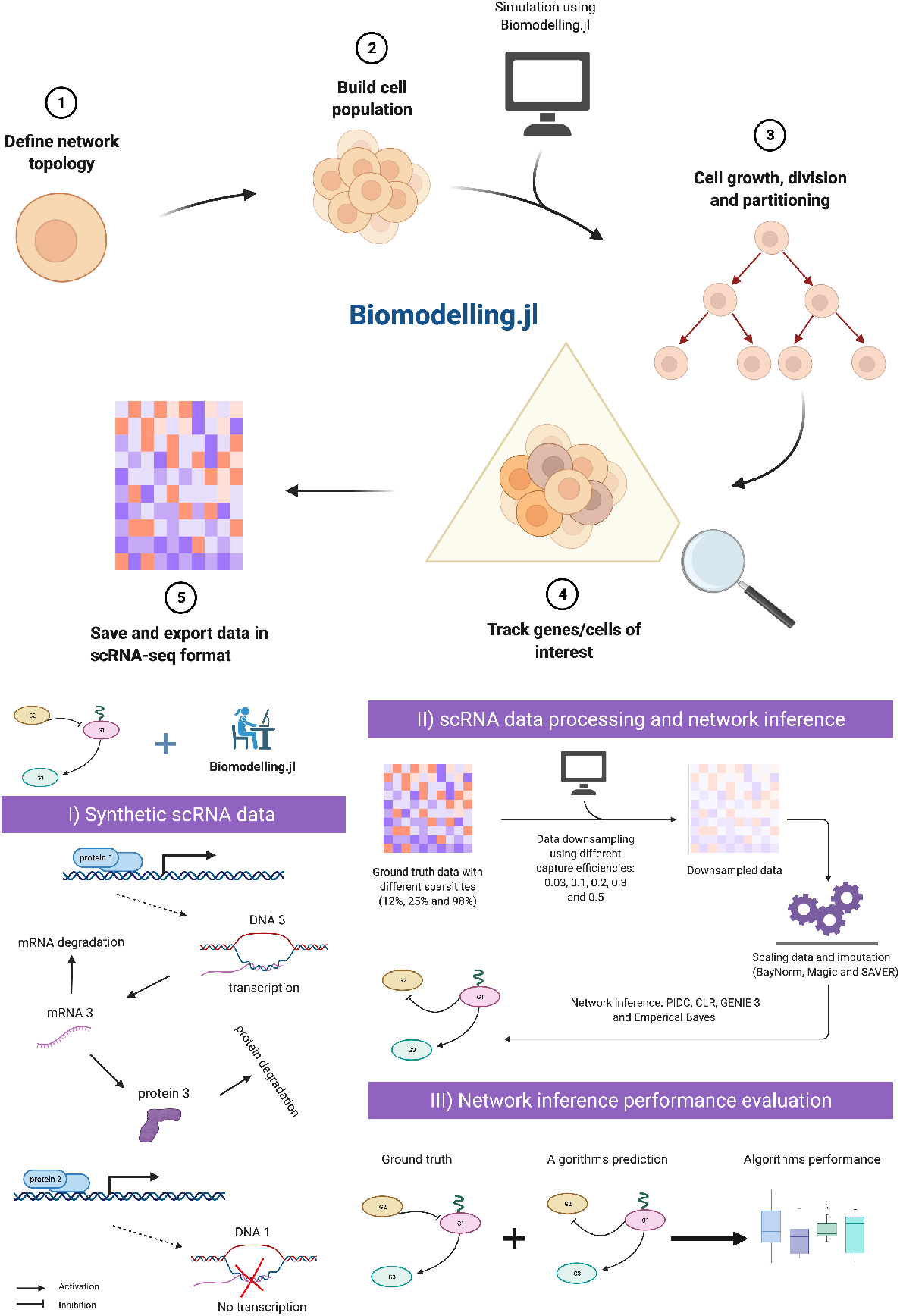
Biomodelling.jl workflow: (1) defining the gene regulatory network topology, interactions and parameters related to gene expression (2) choosing the number of cells and parameters related to cell population such as cell volume control and division noise, (3) couple a stochastic simulation algorithm of biochemical reactions with cell growth, size and division and simulate the cell population, (4) track genes or cells of interest and finally (5) save and export the data in a matrix similar to scRNA-seq data format. (I) synthetic data are generated using *Biomodelling.jl* using different sparsities as described in the Methods section 2.1.1, (II) the obtained data are downsampled, then imputation and network inference are performed as described in Methods section 2.3 and 2.4, finally (III) network inference algorithms predictions are compared with the GT network using metrics presented in Methods section 2.5.

#### 2.1.1 Network topology, sparsity and simulation

In this study, we considered two different types of topology. The first one consists of random connections allowing genes to be regulated by at most one other gene. This topology is referred to in the manuscript as random one regulation (ROR). The second topology considered in this study is a scale free (SF) network topology [44]. Growing evidence has suggested that gene regulatory networks follow a scale free topology [45, 46]. The function *static_scale_free()* from *LightsGraphs* Julia Package (v1.3.5) was used to generate SF topologies. Introducing this more realistic topology means that genes may be regulated by multiple other genes; we allowed for at most four regulators for each gene. In this study, 20-gene and 50-gene regulatory networks were considered.

GRNs are known to be sparse [47, 48, 49] and characterised by a relatively small fraction of regulatory links between genes. In order to evaluate the effect of network sparsity on the performance of inference methods, we considered different levels of sparsity in the simulated networks defined as percentages of all possible links in the GRN excluding self-regulation. Specifically, we used sparsities corresponding to 2.5%, 5% and 10% of possible connections for 20-genes network and 1%, 2% and 4% for 50-genes network. We note that by choosing these sparsity levels we make sure that the percentage of possible connections is kept the same for both networks. Though the graphs generated were directed, in this study we only used undirected information since the network inference methods only outputted this information (apart from GENIE3). As an example, 2.5% of possible links in a 20-gene network corresponds to 5 links that were simulated using the two topologies mentioned in the previous paragraph.

Several chemical reactions stochastic simulation methods have been implemented in *Biomodelling.jl*, the stochastic simulation algorithm (SSA), tau leaping, adaptive tau leaping and non negative poisson tau leaping [50, 51, 52]. For the purpose of this paper, only tau leaping or SSA have been used to simulate the chemical reactions. Our single cell level model simulates gene transcription at a rate which depends on the cell volume, with the transcription rate of a gene in cell *i* being

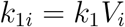

where *V_i_* is the volume of cell *i* and *k*_1_ is the basal transcription rate. This kind of transcription scaling has been reported in mammalian and yeast cells [43, 53, 8], where the authors showed that the numbers of constitutive and inducible mRNAs scale with cell size. We also simulate translation, mRNA decay, protein decay, activation and inhibition as shown in Figure 1 (I).

#### 2.1.2 Types of reactions simulated

Activation and inhibition reactions were modelled as Hill functions *f_act_* and *f_inh_* respectively and defined as follows for a given activator/inhibitor X

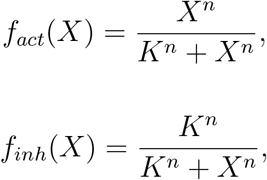

with *n* represents the Hill coefficient and *K* being the microscopic dissociation constant. If gene Y is activated or inhibited by gene X its transcription rate becomes *k*_1*i*_ = *k*_1_*V_i_f_i_*(*X*) for *i* = *act* or *inh*. In the case where a gene X is regulated by multiple genes, we considered two scenarios, the first one is independent or additive (where we sum the regulators’ Hill functions) and the second scenario is synergistic or multiplicative (where we take the product of the regulators’ Hill functions). By allowing a gene to have multiple regulators, we considered three types of *combination reactions* which we refer to as combined activation, combined inhibition and combined action. Combined activation refers to the case where all regulators are activating the gene, combined inhibition refers to the case where all regulators are inhibiting the gene and combined action refers to the case where some of the regulators activate the gene and some of them inhibit the gene.

For example, if Y activates X and Z inhibits X then the transcription rate of X becomes in the multiplicative case (multiplicative combined action)

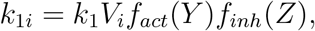

or can be written for the additive case (additive combined action) as follow

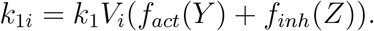

### 2.2 Parameters for mammalian cells

In [54], the authors simultaneously measured absolute mRNA and protein abundance and turnover by parallel metabolic pulse labelling for more than 5000 genes in mammalian cells and reported data for protein and mRNA numbers as well as halflives, transcription and translation rates. To select realistic parameters for accurate GRN simulations, we fitted multivariate Log Normal distributions to data extracted from the aforementioned study using maximum likelihood estimation technique and presented the results in Figure 2. Samples of Protein decay, transcription and translation rates are presented in Figure 2 panels (B), (C) and (D) respectively. We found little correlation between any of the parameters and that the marginal distributions are positively skewed meaning that the majority of the data consists of lower values and the majority of outliers are higher values. To avoid computations taking too long, we also excluded parameter sets that resulted in protein numbers greater than 100,000.

**Figure 2:**
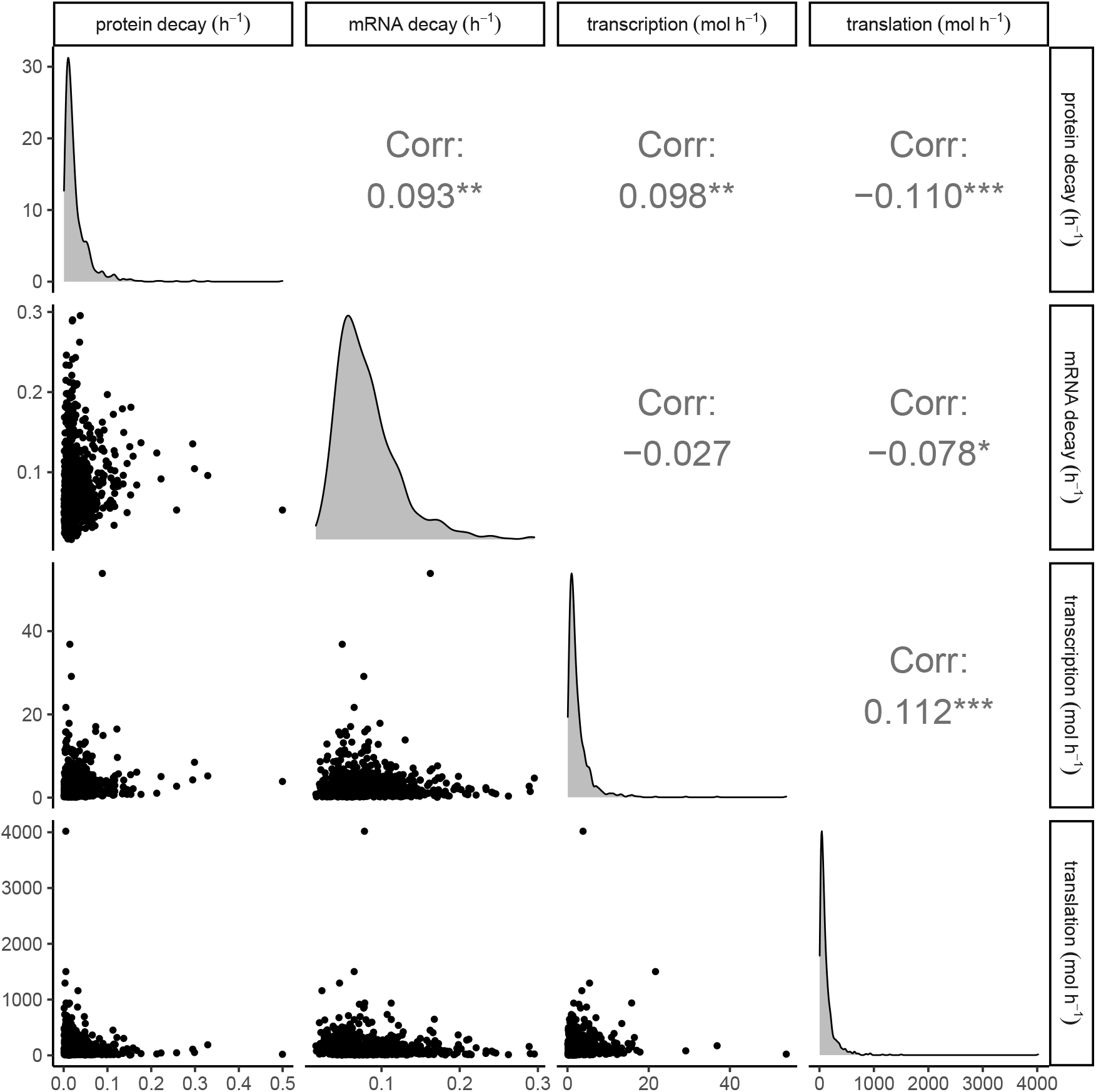
Density plots, scatter plots and correlations of 1000 parameter sets sampled from a multivariate normal distribution fitted to experimental data [54]. Diagonals show distributions of protein decay rates, mRNA decay rates, transcription and translation rates respectively. Lower left scatter plots show relationships between parameter values and upper right plots show Pearson correlation values.

Furthermore, we constrained the choice of the remaining parameters to be realistic and in accordance with experiments. Cell numbers were uniformly sampled from [2, 3] which is consistent with typical scRNA-seq experiments [55]. We note that breakthroughs in technology have allowed even higher numbers of cells to be studied [56]. The cell growth rate was fixed to correspond to a 50 hours doubling time, though we note that we tried a range of doubling times between 24 and 50 hours, which is consistent with mammalian cell doubling times but did not find any consequence for network inference performance. The Hill coefficient *n* was sampled from a log uniform distribution with lower bound 1 and upper bound 10 and the microscopic dissociation constant *K* was chosen to be proportional to the mean value of the steady-state of the regulator in absence of regulation. Finally, we note that the exponent of the power-law degree distribution was sampled from the uniform distribution with bounds [2, 3], which is consistent with [57]. For reproducibility purposes, a list of the 100 parameter sets used can be found here* ^2^.

#### 2.2.1 Cell population: growth, division and partitioning

Without loss of generality, cells were assumed to grow from approximately *V* = 1 at birth to *V* = 2 at division with cell growth rates chosen to correspond to biologically feasible doubling times as explained above. Cell growth was modelled to be exponential

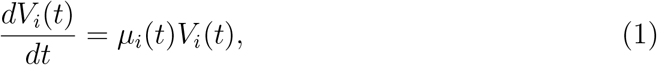

where *μ_i_*(*t*) is the growth rate at time *t* in cell *i*.

To model division noise we adopt the approach of [58, 59] where the final volume of the cell at generation *n* was found to follow a noisy linear map, i.e., the final volume *V_F_* of a given cell was assumed to follow

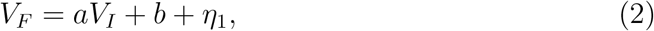

where *V_I_* is the initial volume of the cell, *a* and *b* are linear function parameters, we note that *a* and b have the same value for all cells, and *η*_1_ is the final volume noise. The value of parameter *a* defines the size control strategy of the cell. It is known that many cell types, including mammalian cells show a so-called adder behavior giving a value of *a* =1 [60]. For simplicity, *η*_1_ was set to 0 in this study. Given the value of *a* and the birth size of about *V* = 1, the value of b is also set to be 1.

A dividing cell of volume *V_F_* is assumed to divide into two daughter cells with volumes *V*_*I*_1__ and *V*_*I*_2__ defined by

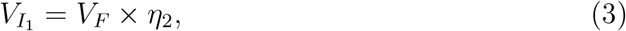

and

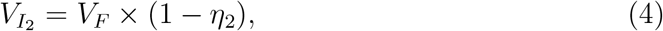

where *η*_2_ represents division noise and is sampled from 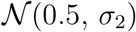. We assumed the contents of the cell are binomially distributed (using *η*_2_) between daughter cells upon division [61]. We note that *η*_1_ and *η*_2_ embed both intracellular stochastic phenomena and also the stochastic influence of extracellular signals. As in [62, 63, 64], in order to keep the population size capped, after a cell division event the new offspring displaces another cell in the population picked at random. Simulating a capped sized population is computationally cheaper than simulating a growing population and leads to more accurate results than using an isolated lineage based approach [65].

To couple the reactions with the exponential growth equation, we ran the stochastic simulation algorithm for a fixed time step before updating the volumes of cells and checking for cell division. This was typically set to *τ* = 0.1 h but we note that we tried smaller time steps as far as *τ* = 0.01 h and found no observable consequences on the simulation output.

#### 2.2.2 Genes tracking and ground truth data

Following the modelling approach described above, genes in the regulatory network were tracked for a given simulation time and data were saved in typical scRNA-seq format (where rows represent genes and columns represent cells). We refer to these data as ground truth (GT) data. In addition, our modelling approach does not only simulate gene expression, it also tracks protein levels in a single cell and stores cell volumes (which are used in data scaling).

### 2.3 Downsampling, scaling and imputation

Given a GT data set and in order to mimic scRNA-seq experiment, as in [19, 66] we assume that the number of transcripts observed in a cell *j* follows a Binomial model with probability *β_j_* (the cell’s specific capture efficiency), which represents the probability of original transcripts in a cell being captured by the sequencing method [66]. In order to simulate downsampling of GT data, the cells’ specific capture efficiencies were obtained from a log-normal distribution centred in *β*, where *β* ∈ {0.03, 0.1, 0.2, 0.3, 0.5}, with a variance set to 0.2, this is consistent with values reported in [67]. The downsampled data from a given capture efficiency *β* is referred to as noisy data (ND-*β*).

In order to perform data scaling, we define the scaling factor (*θ*) for a cell *i* as follows

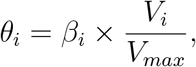

where *V_i_* is cell *i* volume, *V_max_* is the maximum volume in the cell population and *β_i_* is cell i capture efficiency. The scaled data (SD-*β*) are obtained by dividing the noisy data by the cell’s specific scaling factor. Our scaling approach is similar to a global-scaling normalisation strategy, where the expected value of the read count for a gene in a cell is proportional to a gene specific expression level and a cell specific scaling factor [68]. The cell specific scaling factor in the data will be proportional to the cell size and cell specific capture efficiency, which motivates the form chosen for *θ*. In the following we describe, briefly, the imputation methods that are considered in this study.

bayNorm [19] is a Bayesian approach to perform imputation. bayNorm generates for each gene in each cell a posterior distribution of original expression counts, given the observed scRNA-seq read count for that gene and the cell specific capture efficiency assuming a binomial model for transcript capture in the RNA-seq process. The resulting posterior distribution of the original counts relised on emperical based method of estimating a prior on each gene by pulling information across all cells. To perform imputation on ND-*β*, we used *bayNorm()* function from bayNorm R package (v1.6.0). The output data are referred to as BD-*β*.

MAGIC [18] shares information across similar cells, via data diffusion, to fill in missing transcripts. This is achieved in four steps: (i) building a nearest neighbor graph based on cell–cell expression distance, (ii) defining an affinity matrix by applying a Gaussian kernel on the principal components of the graph, (iii) applying a diffusion process on the similarity matrix to obtain a smoothed affinity matrix, (vi) computing the new expression of each gene as a linear combination of the same expression in similar cells, weighted by the similarity strength obtained in the previous steps. To perform imputation on ND-*β*, we used *magic()* function from Rmagic R package (v2.0.3). The output data are referred to as MD-*β*.

SAVER [14] pools information across genes and cells to provide accurate expression estimates for all genes and impute the missing values. SAVER assumes that the count of each gene in each cell follows a Poisson–gamma distribution mixture. The Poisson distribution approximates the technical noise, whereas the uncertainty in the true expression is modelled as a gamma distribution. The recovered expression is a weighted average of the normalized observed counts and the predicted true counts. To perform imputation on ND-*β*, we used *saver()* function from SAVER R package (v1.1.2). The output data are referred to as SAD-*β*.

We refer the reader to [69], a recently published review and benchmarking study that assesses performance, the code quality and the computational time for the above mentioned methods.

### 2.4 Network inference algorithms

We consider four different methods: Information Measurement (PIDC) [9], Emperical Bayes (EB) [70], Context Likelihood of Relatedness (CLR) [9], and GENIE3 [71], see Figure 1(II). The overall workflow of the aforementioned methods focuses on modelling the relationship between genes using different correlation metrics.

PIDC and EB were developed by the same authors with EB presented as an improvement of PIDC. Both methods use partial information decomposition (PID) as follows: (i) compute the mutual information between two genes X and Y and the unique mutual information between X and Y given a third gene Z, (ii) define the proportional unique contribution (PUC) between two genes X and Y as the sum of the ratio of unique to mutual information calculated using every other gene Z in a network, (iii) an empirical probability distribution is estimated from the PUC scores for each gene, and the confidence of an edge between a pair of genes is given. EB provides an additional step to smooth the empirical distributions using a regression-based mode-matching method. The methods output a ranked list of undirected edges using the confidence scores obtained. The Julia implementation of these methods was used: *InformationMeasures.jl* (v0.3.1), *NetworkInference.jl* (v0.1.1) and *EmpiricalBayes.jl*

CLR computes the mutual information between two genes and calculates the statistical likelihood of each mutual information value within its network context. Then, the pairwise genes mutual information is compared to the background distribution of mutual information scores for all possible gene pairs. The most probable interactions are those whose mutual information scores stand significantly above the background distribution of mutual information scores. The Julia implementation of this method in the following packages *InformationMeasures.jl* (v0.3.1) and *NetworkInference.jl* (v0.1.1) was used.

Originally developed for bulk RNA-seq and best performer in the Dialogue for Reverse Engineering Assessments and Methods (DREAM4) challenge, GENIE3 is widely applied to scRNA-seq. Unlike many methods in the same category that look at gene pairs or gene triplets, GENIE3 takes into account the interaction of an arbitrary number of genes in one calculation and can capture the nonlinear dependencies between genes by decomposing the prediction of a regulatory network between p genes into p different regression problems. Although GENIE3 can return a directed network, for the sake of comparison with the other methods, we considered the undirected network option. We used *GENIE()* function from GENIE3 R package (v1.10.0).

As control we also report random inference (RAND), which returns for a given sparsity random links in the GRN. We note that in this systematic study we matched the network’s sparsity to the inference method algorithms’ threshold, meaning that if for a given sparsity the GT network has N links, we chose the inference algorithms’ threshold that returns the top N predicted links.

We refer the reader to [11], a recently published review that assesses the code implementation and usability and the computational time of the above mentioned methods, with the exception of CLR.

### 2.5 Network inference performance evaluation

To evaluate the network inference algorithms performance, we consider two metrics: Area Under Receiver Operating Characteristic curve (AUROC) [72] and Area Under Precision-Recall curve (AUPR) [73], see Figure 1(III).

The ROC curve is defined as a plot of False Positive Rate (FPR) versus True Positive Rate (TPR) (also known as sensitivity or recall) which are given in function of True Positive (TP), True Negative (TN), False Positive (FP) and False Negative (FN) as follow

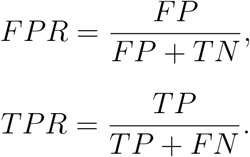

The AUROC is then easily obtained from the ROC curve, many options are available, we used *AUC()* function from DescTools R package (v0.99.39) that takes as input the ROC curve and the method to compute the area, we chose ‘*trapezoid*’. AUROC is characterised by the absence of bias toward models that perform well on the minority class at the expense of the majority class, in other words AUROC does not favour methods that are good at identifying interactions between genes while failing to detect the absence of interactions [74].

The PR curve is defined as a plot of TPR against Precision (P) which is given as

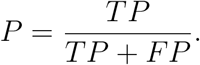

The AUPR is obtained from PR curve using *AUC()* function as described above. Using AUPR we are able to assess the performance of a method on the minority class, in other words, since the gene regulatory networks are sparse, we can assess the performance of a given method on how it does in detecting existing interactions between genes [74].

## 3 Results

### 3.1 Synthetic scRNA-seq data for a toy example: unscaled expression leads to uniformly high and positive correlations

We used the pipeline described in Figure 1 to investigate different scenarios for network inference. We begin in this section by presenting a toy example using our method of synthetic scRNA-seq data generation (Figure 1). This example serves to show typical output of our simulation pipeline and also illustrates the difficulties of performing accurate network inference using scRNA-seq data.

While we only make use of the final time point for mRNA and cell volume in this study (as scRNA-seq is obtained in a time snap-shot), we present plots of the full volume time series for a single cell along with the corresponding levels of mRNA and protein in Figure 3A-C. For initial conditions we chose the steady state mean value of mRNA and protein species in the absence of any regulation. Furthermore, by using only the final time point for network inference, we ensured all simulated cells are uncorrelated from the initial condition. As we made clear in Methods section 2.2 our choice of parameters such as cell doubling time, transcription, translation and decay rates keep the mRNA and protein numbers within biologically feasible levels for mammalian cells. However, we note that our approach can also be adapted for any other cell type by using different parameterisations.

**Figure 3:**
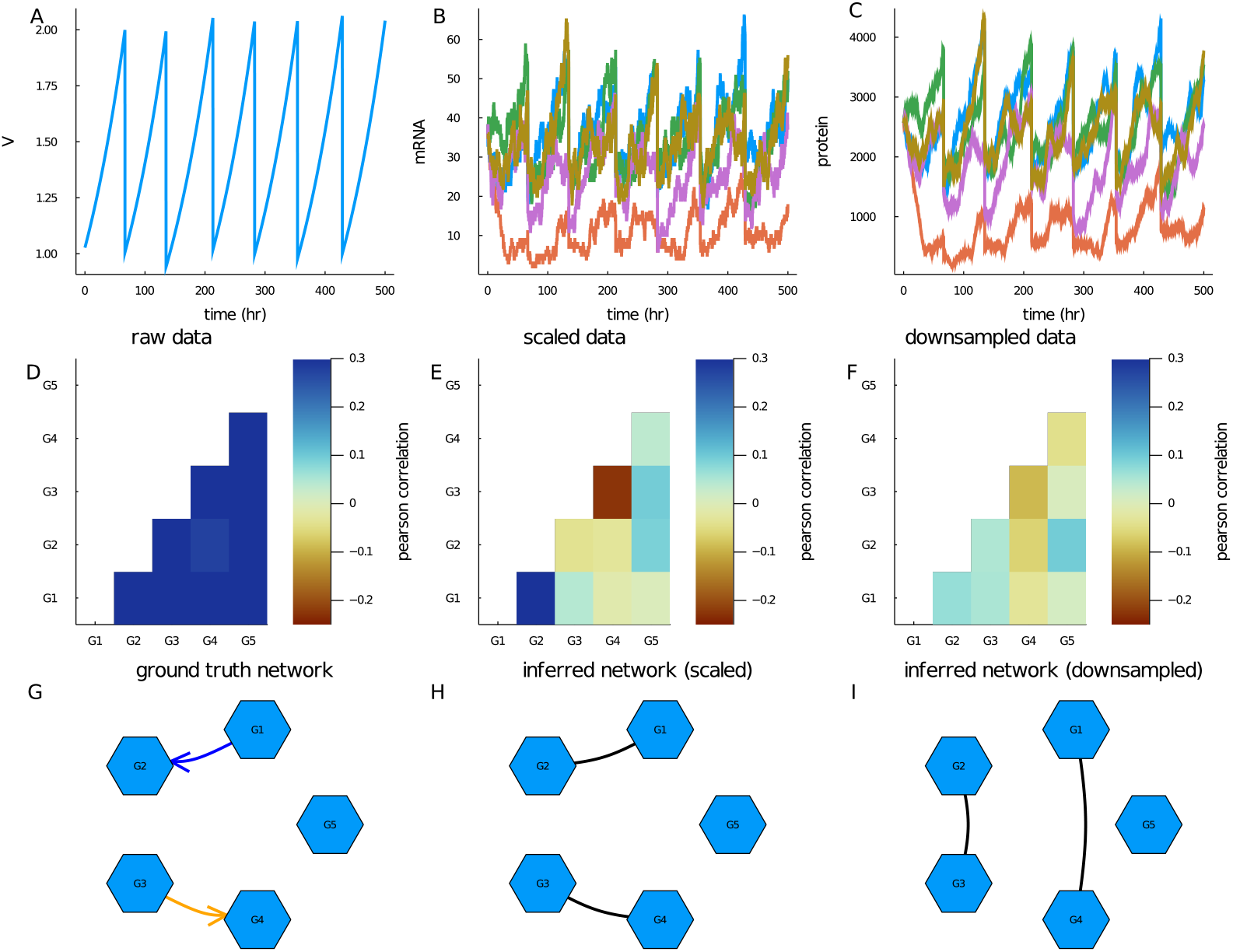
Synthetic scRNA-seq data generated for 5 gene network example. The network was simulated using 500 cells over a 500 hour time period with parameters sampled as described in Methods section 2.2. (A) Plot of the volume time series of a single representative cell. Early divisions are due to replacement in order to keep number of tracked cells constant. (B) Plot of corresponding mRNA time series for the 5 genes modelled. (C) Plot of corresponding protein time series for the 5 genes modelled. (D) Heatmap of mRNA pearson correlations taken from final time point. (E) Heatmap of mRNA pearson correlations scaled by cell volume taken from final time point. (F) Heatmap of mRNA pearson correlations scaled by cell volume and subsequently downsampled using Binomial downsampling with 20% capture efficiency. (G) Graph of ground truth network where a blue arrow represents a link with an activating reaction and an orange arrow represents a link an inhibiting reaction. (H) Graph of inferred reaction network obtained from PIDC algorithm using mRNA data scaled by cell volume at final time point as input. Predicted links are represented by solid black lines. (I) Graph of inferred reaction network obtained from PIDC algorithm using mRNA data scaled by cell volume at final time point and downsampled (using Binomial downsampling with 20% capture efficiency) as input. Predicted links are represented by solid black lines.

In Figure 3D, we show gene-gene correlations computed from the cell population at the final time point across the 5 genes. Strikingly we found that without scaling the raw mRNA copy numbers by cell volume, gene correlations are dominated by cell volume (see Figure 3D). This is because gene expression scales with cell size and therefore mRNA levels for different genes therefore have a global positive correlation due to cell size scaling. Hence any correlations due to activations or inhibitions are obscured by the cells position in the cell cycle. This information can be retrieved by dividing the raw mRNA copy numbers by the cell volumes (as shown in Figure 3E). Inspecting Figure 3E we can observe a strong positive correlation between gene 1 and gene 2 and a strong negative correlation between gene 3 and gene 4. This is consistent with what we would expect from the ground truth network (illustrated in Figure 3G). While, most scRNA-seq protocols do not measure cell size (see [6] for an exception), one can correct for cell size scaling in real scRNA-seq data by normalising by total transcript counts per cell, which is expected to scale with cell size [6].

Drop-out events are one of the most important features of single cell data. While their technical origin is hotly debated, the evidence for zero-inflation has been questioned as the statistics of drop-out events are consistent with a simple model of binomial capture of original transcripts during scRNA-seq protocols [19]. To investi-age the effect of drop-outs, We next artificially induced drop-out events to the final mRNA data (before scaling by final cell volume). We downsampled our data using a Binomial distribution with capture efficiency of 20%, see Methods section 2.3 for more details. This approach is similar to the method used to generate single cell simulation data for network evaluation that was published recently [75]. As shown in Figure 3F downsampling in this manner removes a significant level of the correlation information.

Finally, we present two network inference results. In Figure 3(H) we show the network inferred using the PIDC algorithm with the final mRNA data divided by final cell volume as input. We selected the threshold parameter to be equal to the sparsity of the network (as we do for the rest of the results presented in this paper). We show in Supplemental Figure 1 that this is the most appropriate parameter choice. By making this choice we focus our study on the impact of imputation on inference accuracy rather than the choice of inference algorithm parameters. We note that in applications to real data, of course the true sparsity will not be known and a best guess should be used. For this simple toy example, we can see that PIDC identified the whole network correctly (comparing Figure 3(H) and (I)). We note that this network is not representative of a real biological network due to its small size. However as shown in Figure 3(I) even in this simple case downsampling the data affects the results significantly and PIDC no longer predicts any correct links. Hence, we observe that downsampling of the data that is associated with low capture efficiency and drop-out in scRNA-seq data represents a challenge for network inference. In the following, we investigate this issue systematically in bigger networks and ask if imputation methods could help to resolve this challenge.

### 3.2 Network inference algorithms tend to perform better for sparser networks

In this section we present the performance of 4 commonly used network inference algorithms using ground truth data (i.e., no downsampling is performed) as input from 100 different simulated networks with 20 genes. Each network was randomly sampled in terms of the links generated, number of cells and parameter values used. For simplicity, we limited the number of links between genes to at most one (i.e., we use a ROR network topology, see Methods section 2.1.1). Though this case is biologically infeasible, we used this to gauge the best case performance of the different algorithms and focus on the impact of network sparsity on network inference. The sparsity parameter relates to the number of links in the network, where a larger parameter leads to more links. We considered network sparsities that correspond to 5, 10 and 19 links present in the network (out of a possible 190). We present the results of commonly used network inference metrics in Figure 4.

**Figure 4:**
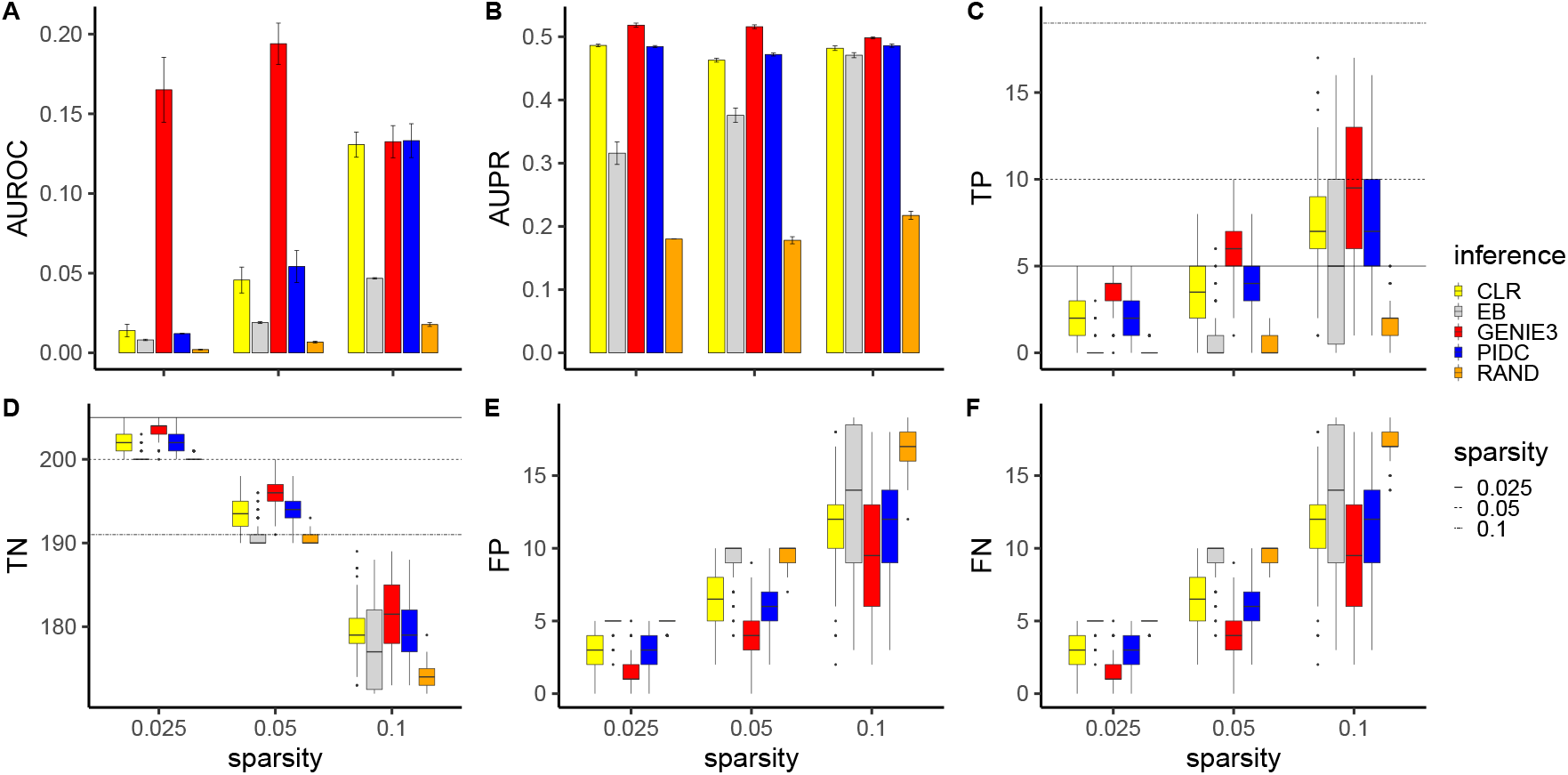
Network inference results using ground truth data (without downsampling) from 100 different simulated random one regulation networks with 20 genes for 3 different network sparsities. Each network was simulated over 500 hours using parameters sampled as described in Methods section 2.2. (A) shows a barplot of the AUROC score for the 4 different network inference algorithms considered as well as a random classifier (RAND). (B) shows a barplot of the AUPR score for the 4 different network inference algorithms considered as well as a random classifier. Confidence intervals for barplots were computed by subsampling 35 out of 100 networks 100 times. (C) shows a boxplot of the true positives found for each network inference algorithm and random classifier for 3 different sparsity levels. The horizontal lines depict the actual number of true positives for reference. (D) shows a boxplot of the true negatives found for each network inference algorithm and random classifier for 3 different sparsity levels. Again, the horizontal lines depict the actual number of true negatives for reference. (E) shows a boxplot of the false positives found for each network inference algorithm and random classifier for 3 different sparsity levels. (F) shows a boxplot of the false negatives found for each network inference algorithm and random classifier for 3 different sparsity levels.

Our first observation is that in general all 4 network inference algorithms perform significantly better than the random classifier (across all measures considered). In terms of ranking, for this data set, it appears that GENIE3 performs the best, followed by PIDC then CLR and finally Empirical Bayes. This is consistent with other studies where it was found that GENIE3 has the best network inference performance for many different data sets [39].

With respect to the GENIE3 algorithm, we observed no clear relationship between the AUROC score and network sparsity (Figures 4A). Similarly, we see that the AUPR score stays relatively constant with respect to sparsity (Figures 4B). However, we noticed a clear trend regarding the number of true positives (Figures 4C) versus network sparsity. As the sparsity parameter is increased, while the number of true positives increases, the overall fraction of average correctly identified true positives decreases (0.8, 0.6, 0.47 for 0.025, 0.05 and 0.1 sparsities respectively). We found a similar trend for the false positives and false negatives (Figures 4E and F) while the true negatives decrease with increasing sparsity parameter (Figures 4D). We note also that the variance in the number of true positives, true negatives, false positives and false negatives increases with sparsity, implying GENIE3 is less reliable for larger sparsity values.

We next considered the PIDC and CLR algorithms which perform similarly in this case. In contrast to the GENIE3 algorithm, we observed an increase in the AUROC score for both these algorithms as the sparsity is increased (Figure 4A). The AUPR score did not change with sparsity (Figure 4B) and the number of true positives increases with sparsity (while the overall fraction of average correctly identified true positives decreases) for both algorithms (Figure 4C). We found a similar trend for the false positives and false negatives (Figures 4E and F) while the true negatives decrease with increasing sparsity parameter (Figures 4D). We note that the CLR algorithm appears to have a constant variance for the number of true positives, true negatives, false positives and false negatives for the different sparsities considered while the same metrics for the PIDC algorithm increases in variance for the highest sparsity.

Unlike the other algorithms considered, Empirical Bayes produces similar trends for both the AUROC and AUPR scores with both increasing with the sparsity parameter. For the lower sparsities considered (0.025, 0.05), the number of true positives, true negatives, false positives and false negatives is similar to the random classifier. However, for the largest sparsity (0.1) the Empirical Bayes algorithm improves upon the random classifier but with very large variance.

### 3.3 Scale-free topologies are challenging for accurate net work inference

Here we build on the previous sections by considering realistic scale-free topologies.

In this case, since more than one link can be made between genes (we allow up to 4 genes to activate/inhibit another gene) using scale-free topologies, we must consider how this regulation occurs. To explore this, we considered two different kinds of regulation, multiplicative or additive (for details see Methods section 2.1.1). We present the results of multiplicative versus additive regulation in Figure 5.

**Figure 5:**
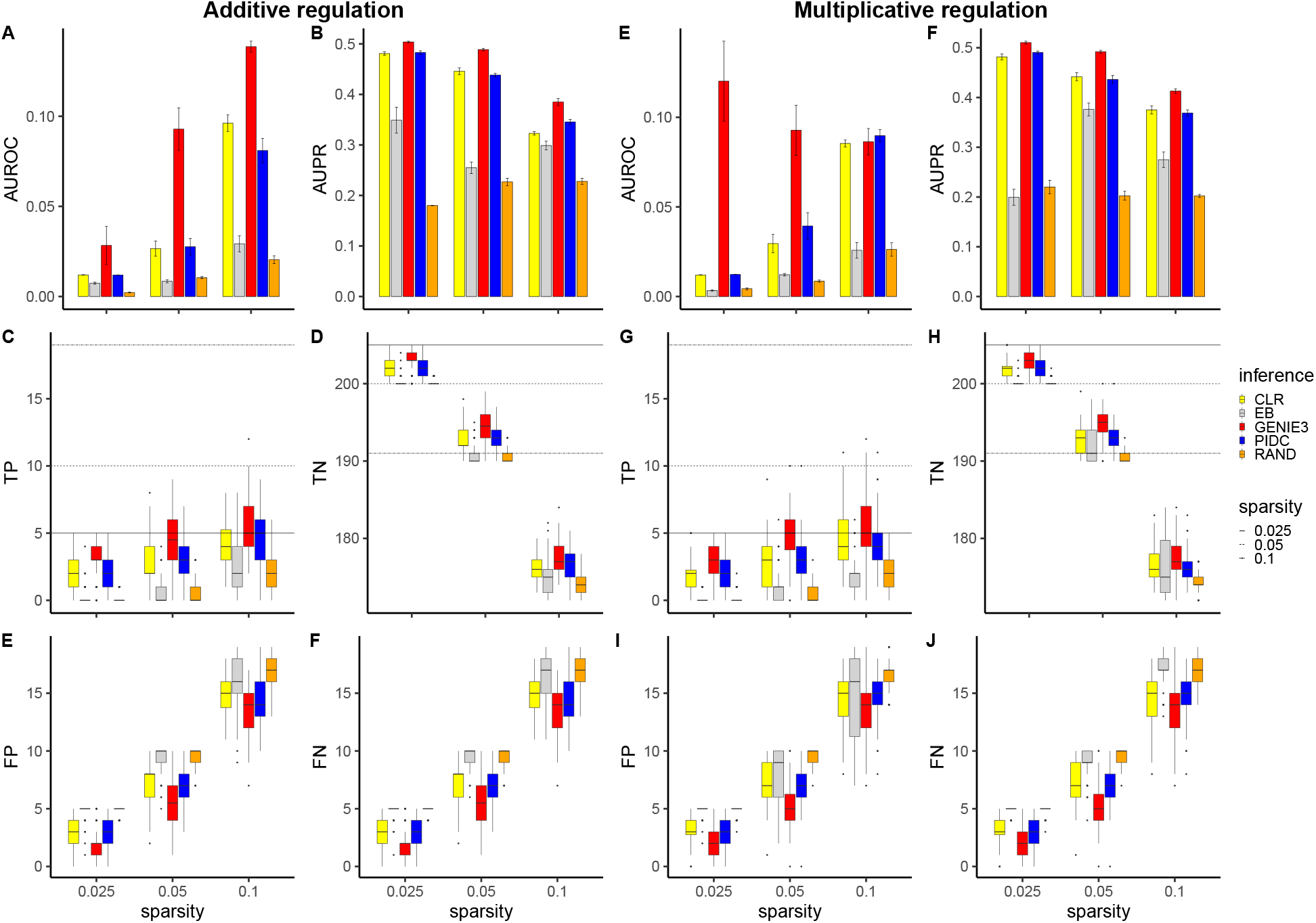
Network inference results using ground truth data (without downsampling) from 100 different simulated scale-free networks with 20 genes for 3 different network sparsities using additive or multiplicative regulation. Each network was simulated over 500 hours using parameters sampled as described in Methods section 2.2. (A) and (E) show barplots of the AUROC score for the 4 different network inference algorithms considered as well as a random classifier (RAND) for additive and multiplicative regulation respectively. (B) and (F) show barplots of the AUPR score for the 4 different network inference algorithms considered as well as a RAND classifier for additive and multiplicative regulation respectively. Confidence intervals for barplots were computed by subsampling 35 out of 100 networks 100 times. (C) and (G) show boxplots of the true positives found for each network inference algorithm and random classifier for 3 different sparsity levels for additive and multiplicative regulation respectively. The horizontal lines depict the actual number of true positives for reference. (D) and (H) show boxplots of the true negatives found for each network inference algorithm and random classifier for 3 different sparsity levels for additive and multiplicative regulation respectively. Again, the horizontal lines depict the actual number of true negatives for reference. (E) and (I) show boxplots of the false positives found for each network inference algorithm and random classifier for 3 different sparsity levels for additive and multiplicative regulation respectively. (F) and (J) show boxplots of the false negatives found for each network inference algorithm and random classifier for 3 different sparsity levels for additive and multiplicative regulation respectively.

Overall, we found the performance is poorer compared to the ROR network topologies results presented in Figure 4, i.e., the results were closer to the random classifier for all algorithms considered. This is due to the scale-free nature of the networks considered as we found very little difference in the performance of the networks produced using additive versus multiplicative regulation. Both forms of regulation display the inverse relationship between the network sparsity parameter and accuracy that we observed in the previous section. This inverse relationship is also reflected in the AUPR scores in Figures 5B and F. Interestingly, the AUROC scores show an opposite trend for additive and multiplicative regulation, with the AUROC score increasing for higher sparsities (apart from the GENIE3 algorithm for multiplicative regulation). We also highlight that the overall ranking of the network inference algorithms were preserved from the ROR case, with GENIE3 again performing the best, followed by PIDC, CLR then Empirical Bayes (which is only slightly better than random classification). While the overall accuracy is diminished from the ROR case, the results appear more robust (i.e., the variance is decreased).

Due to inconsistencies we observed using the common AUROC and AUPR scores, we use an easier to interpret score, the precision, for the remainder of the paper. Since we fix the threshold used in the network inference algorithms to the sparsity of the network, the precision can be interpreted simply as the fraction of correctly identified true positives.

### 3.4 Different imputation methods perform better for different network inference methods

To address the question of which imputation method is best for the purpose of accurate network inference we generated synthetic scRNA-seq data for 100 scale-free network topologies using 20 genes. For simplicity, we only present results for the middle sparsity case from previous sections (i.e., sparsity = 0.05 or 10 out of 190 possible reactions have links) and use multiplicative regulation (since both additive and multiplicative regulation gave similar results). To reflect real scRNA-seq data, we downsampled our data using capture efficiencies that reflect current technologically possible average capture efficiencies [76]. We present the results as boxplots in Figure 6 where the first row corresponds to the precision scores for different network inference algorithms using different imputation methods.

**Figure 6:**
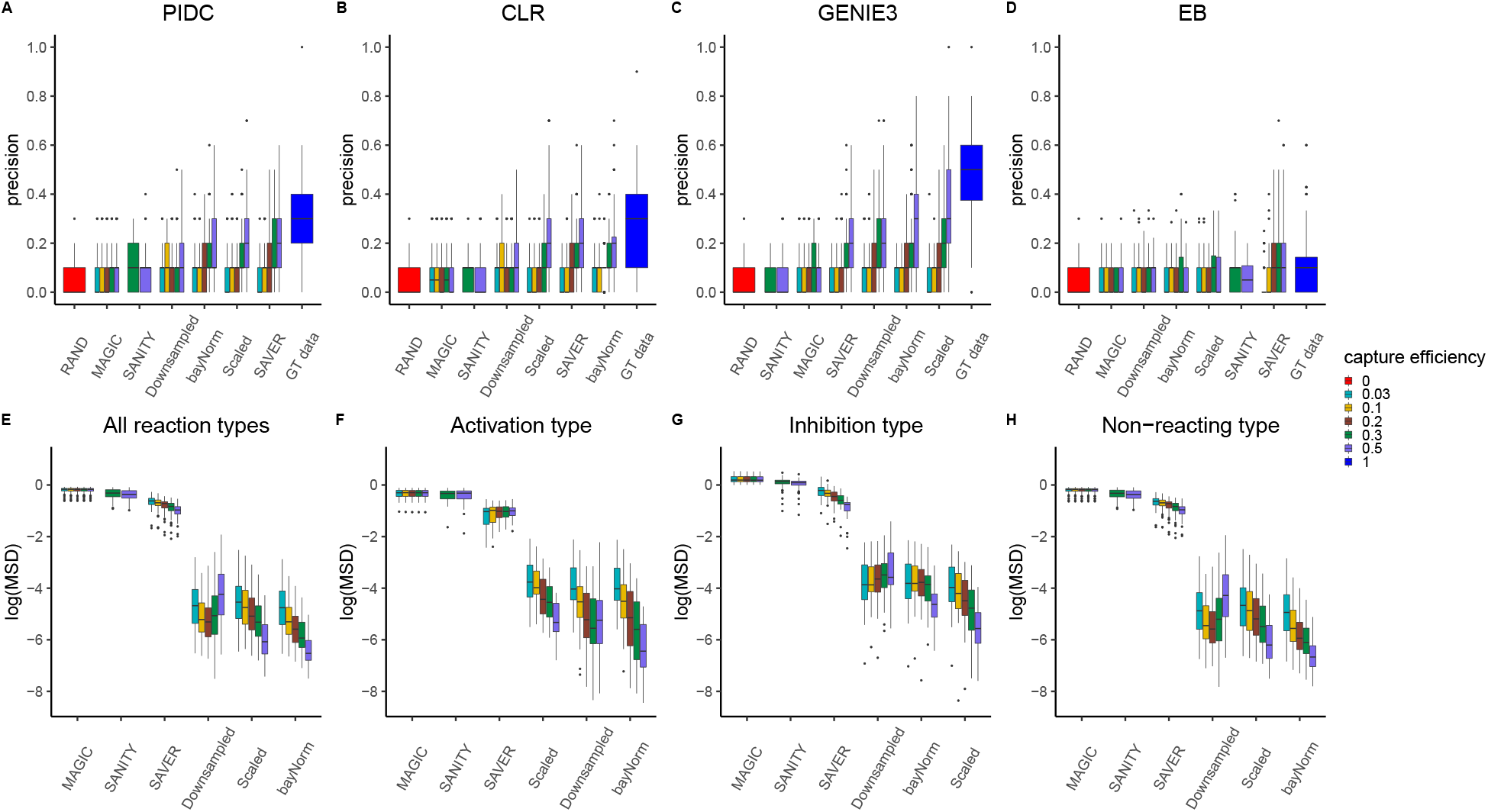
Impact of imputation on network inference performance and gene-gene correlation preservation for 100 different simulated 20 gene networks using sparsity = 0.05 with multiplicative regulation for various capture efficiencies. Figures (A) to (D) show boxplots of precision scores obtained for different imputation algorithms displayed on x-axes for PIDC, CLR, GENIE3 and Empirical Bayes respectively. RAND corresponds to precision obtained using random classification and GT data corresponds to precision obtained without downsampling (i.e., capture efficiency is set to 1). Figures (E) to (H) show the mean squared deviation between gene-gene correlations obtained using the ground truth data and those obtained using various imputation methods displayed on x-axes (with results plotted on a log-scale). Figure (E) show results obtained using all reaction types, while Figure (F), (G) and (H) show results obtained using only activation, inhibition and non-reacting type reactions respectively.

Overall we found that no imputation method is able to completely recapitulate the network inference results obtained using the ground truth data. There is also a general trend where as the capture efficiency decreases, the performance of the network inference decreases, with no network inference method/imputation method combination improving upon random classification for capture efficiencies less than 10%. Another general trend we notice is that MAGIC and SANITY imputation methods lead to very poor network inference accuracy for all network inference methods studied and all capture efficiencies. We also note that the SANITY imputation algorithm failed to converge for capture efficiencies lower than 30%. We also highlight that we ordered the results on the x-axes by average score and highlight that ‘downsampled’ corresponds to no imputation performed, hence every method to the right of ‘downsampled’ is beneficial for inference.

Inspecting each individual network inference algorithm, we found that no one imputation method works best for every network inference algorithm. In Figure 6A we observe that the SAVER imputation method works best on average when combined with the PIDC algorithm with scaled data and bayNorm performing very similarly. For the CLR algorithm, bayNorm performs best on average, followed closely by SAVER (see Figure 6B). For GENIE3 which produces the best ground truth performance, scaled data gives the best results followed by bayNorm (see Figure 6C). Remarkably for the 50% capture efficiency scaled case, there is a single network which is inferred exactly. Figure 6D shows the Empirical Bayes algorithm results which works best when combined with SAVER imputation, though it should be noted that even for ground truth data Empirical Bayes performance is only marginally better than random classification.

We next investigated how well different imputation methods preserved gene-gene correlations. To do this we first computed gene-gene Pearson correlations in the ground truth data for the 100 synthetic scRNA-seq data sets. We then computed the corresponding Pearson correlations for various imputation methods for different capture efficiencies. We show one such example for each imputation method and for three different capture efficiencies in Supplemental Figure 2. From this figure we see a general trend where the gene-gene correlations become less correlated with the ground truth data as the capture efficiency was decreased. We can also notice a pattern emerging with inhibition reactions (highlighted in grey) being less well preserved than other reaction types. We also noticed that the SAVER imputation method seemed to artificially inflate correlations. To test these observations more robustly, we computed the mean squared deviation between gene-gene correlations obtained using the ground truth data and those obtained using various imputation methods for all 100 data sets. We present these mean squared deviations as boxplots in the second row of Figure 6.

In general, the bayNorm imputation method preserved the gene-gene correlations best (see Figure 6E). The only other method improving on ‘downsampled’ was the scaled method which performed similarly well. MAGIC, SANITY and SAVER performed poorly in preserving gene-gene correlations, however SAVER appeared to improve with increasing capture efficiency. In Figure 6F we show the mean squared deviations found using only activation type reactions, and here we found that only bayNorm improves over the gene-gene correlations found using the downsampled data. We also observed that SAVER and MAGIC do not improve in performance with increasing capture efficiency for activation type reactions. In Figure 6G, we show that the scaled method performed best at preserving gene-gene correlations of inhibition type reactions, followed closely by bayNorm which also improved upon the downsampled data. Finally we observed that bayNorm is best at preserving gene-gene correlations for non-reactions (see Figure 6H).

### 3.5 Overall performance of network inference algorithms is inversely related to number of combination reactions considered

To examine the impact of the number of genes on overall network inference performance, in this section we extended the size of the networks analysed from 20 to 50 gene networks. We used sparsities such that the fracton of links present in the network were consistent with the 20 gene case from previous sections. This also prevented the maximum degree of the network from exceeding the maximum of 4 which is currently supported in Biomodelling.jl. We present the results for sparsity = 0.02 as boxplots in Figure 7, as in the previous section, where the first row corresponds to the precision scores for different network inference algorithms using different imputation methods.

**Figure 7:**
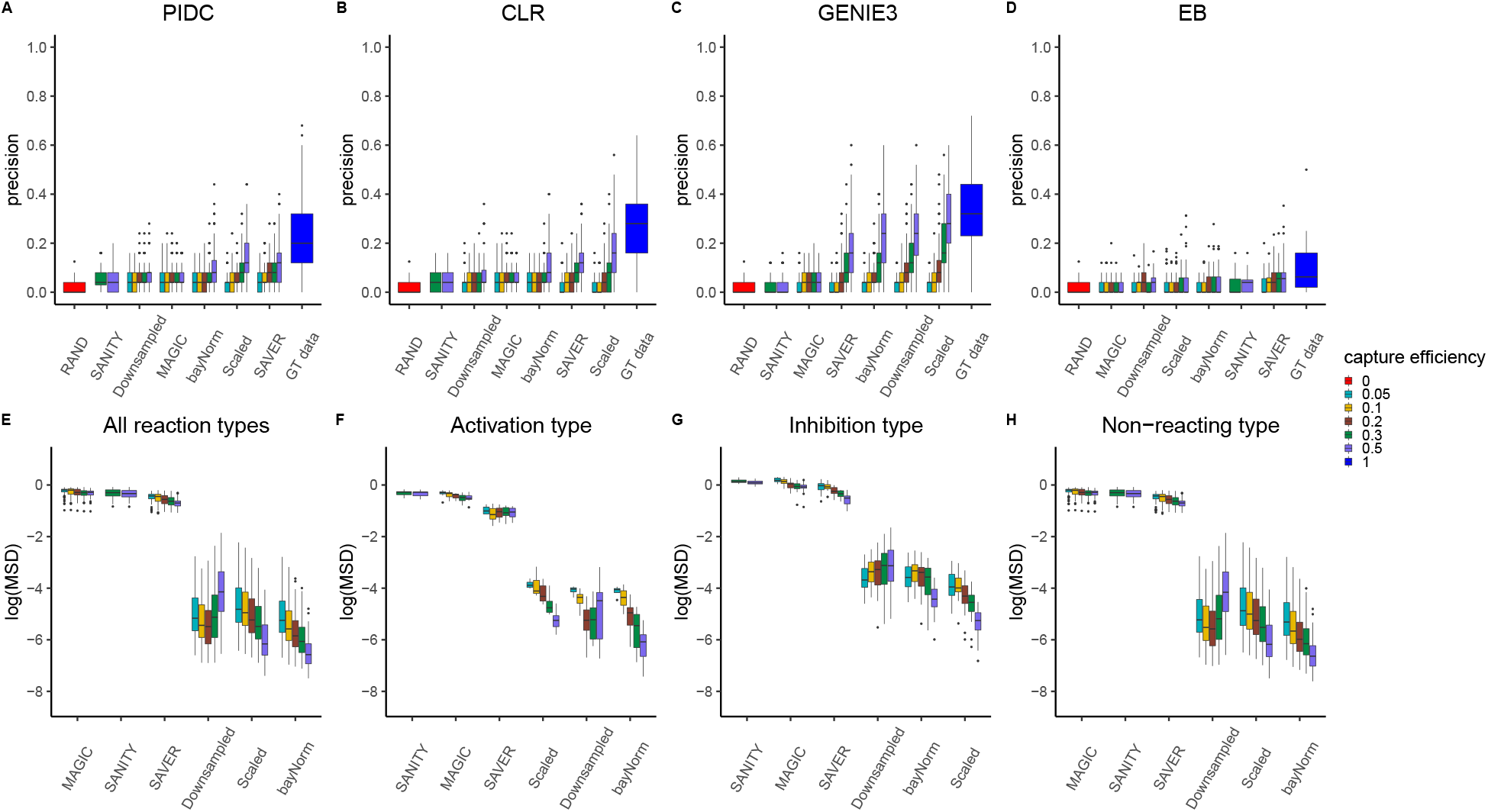
Impact of imputation on network inference performance and gene-gene correlation preservation for 100 different simulated 50 gene networks using sparsity = 0.02 with multiplicative regulation for various capture efficiencies. Figures (A) to (D) show boxplots of precision scores obtained for different imputation algorithms displayed on x-axes for PIDC, CLR, GENIE3 and Empirical Bayes respectively. RAND corresponds to precision obtained using random classification and GT data corresponds to precision obtained without downsampling (i.e., capture efficiency is set to 1). Figures (E) to (H) show the mean squared deviation between gene-gene correlations obtained using the ground truth data and those obtained using various imputation methods displayed on x-axes (with results plotted on a log-scale). Figure (E) show results obtained using all reaction types, while Figure (F), (G) and (H) show results obtained using only activation, inhibition and non-reacting type reactions respectively.

We found a general deterioration in the performance of all network inference algorithms with the medium sparsity for the 50 gene case performing worse than medium sparsity for the 20 gene case (compare Figure 7A to D with (Figure 6A to D). While GENIE3 still performed the best overall, CLR performed better than PIDC in this case. Empirical Bayes was found again to be only marginally better than random classification. In terms of imputation methods, we found that SAVER worked best when combined with PIDC or Empirical Bayes and the scaled method worked best for CLR or GENIE3 algorithms. This is broadly consistent with the 20 gene case.

Surprisingly, the gene-gene correlations are very closely aligned with the 20 gene case (compare Figure 7E to H with Figure 6E to H), even for different reaction types. This implies that the source of the deterioration in network inference is elsewhere. To further investigate this, we also examined the number of combination reactions (see Methods section 2.1.1 for details). We present the number of combination reactions for the 20 gene and 50 gene cases in Supplemental Figure 6. We found a significant increase in the total number of combination reactions in the 50 gene case versus the 20 gene case. Furthermore, we found that if we approximately matched the number of combination reactions for different number of genes (e.g. 0.02 sparsity for the 50 gene case and 0.1 sparsity for the 20 gene case) we observed a very similar performance (compare Supplemental Figure 7). Hence, this implies that it is not the gene number that dictates the overall performance but the number of combination reactions present in the network.

## 4 Discussion

Here we have introduced Biomodelling.jl, an open source julia package, for producing synthetic scRNA-seq datasets based on a known gene-regulatory network. Biomod-elling.jl simulates realistic stochastic gene expression coupled to cell size in growing and dividing population of cells using an agent-based approach. Downsampling using a binomial distribution is used to model capture efficiency and drop-out in scRNA-seq protocols. While there are other methods available for generating synthetic scRNA-seq datasets such as GeneNetWeaver and Splatter, these do not account for gene-gene correlations that arise due to an underlying gene regulatory network and cell growth. Hence, Biomodelling.jl can be used for benchmarking network inference methods. In this study, we investigated the effectiveness of imputation on recovering gene-gene correlations that are lost due to drop-out.

We first demonstrated the use of Biomodelling.jl by presenting results from a toy 5 gene network example. This showed that to uncover true gene-gene correlations it was necessary to scale the raw mRNA numbers by cell volume, otherwise gene-gene correlations would be uniformly high and positive. Without scaling by cell volume, the mRNA numbers per cell for each gene are dominated by their position in the cell cycle. While, there are several methods that have been developed to remove cell cycle effects for scRNA-seq studies [77, 78, 79], we propose for the purpose of removing cell size effects one could use a total count normalisation. We matched the threshold parameter of the network inference algorithms with the sparsity of the network, as this yields the best performance and simplifies the interpretation of the performance. For this simple network, PIDC was able to correctly identify the whole network if the volume scaled mRNA data was used. However we found that drop-out events simulated by downsampling lead to poor network inference performance, implying even for very simple networks imputation may help. We note that in general the sparsity of network is not known, but we suggest the threshold could be derived from the number of known transcription factors present in the considered network.

We then explored the performance of common network inference algorithms for simple topologies (ROR) using 20 genes network topologies. We found that all the network inference algorithms considered performed significantly better than random classification (apart from Empirical Bayes). Furthermore, GENIE3 performed best in this setting and sparser networks were generally easier to infer. Introducing scale-free topologies led to a general deterioration in the performance of the network inference algorithms but the overall ranking of the algorithms was retained from the ROR network topologies case. We also observed very little difference using additive or multiplicative regulation. Hence we decided to use multiplicative scale-free topologies for evaluating the impact of imputation methods on the performance of network inference algorithms.

We next examined the impact of performing imputation prior to applying the network inference algorithms for a range of experimentally feasible capture efficiencies. In general we found that inference performance was inversely related to the capture efficiency regardless of imputation method used and that even for higher capture efficiencies the imputation methods were never able to completely recapitulate the ground truth data case, though they frequently improved upon just using the downsampled data. The best choice of inference algorithm depended on the choice of imputation method, i.e., there was no one best imputation method for every network inference algorithm. Though we found clear evidence that some imputation methods should not be used for network inference. SAVER, bayNorm and the scaled method can be used depending on the choice of inference algorithm, for example SAVER and PIDC worked well together. We found that MAGIC and sanity imputation methods never improved upon using the downsampled data for any network inference algorithms that we considered.

To better understand the network inference results we also examined how well gene-gene correlations were preserved using several imputation methods. Overall, we found that bayNorm was the best at preserving the gene-gene correlations found in the ground truth data. We also examined the gene-gene correlations for specific reaction types. Only bayNorm performed better than down-sampled data for activation type reactions, while bayNorm and the scaled method performed better than downsampled data for inhibition and non-reacting type reactions. The fact that SAVER performs so poorly here is inconsistent with the performance we found for network inference. Therefore we examined this further, and while the gene-gene correlations are in general higher than the ground truth gene-gene correlations, we found that they are off by a constant (approximately the median correlation). In other words, the overall order or ranking of correlations is preserved which may explain why the network inference algorithms such as PIDC worked well with SAVER.

We also examined the impact of increasing the size of the gene network simulated. Across all network inference algorithms, we found a deterioration in the quality of the inference. We also computed the gene-gene correlations for various imputation methods for these larger networks but unexpectedly found no difference compared to the smaller gene networks, implying the source of the deterioration was elsewhere. We found that the performance of the network inference seemed to be proportional to the number of combination reactions (where it is possible to have a gene activated and inhibited simultaneously) with similar performance recorded for 20 gene networks and 50 gene networks with the same number of combination reactions. We speculate that incorporating protein information into inference may help improve performance in such networks.

Finally, we compared our results with two recent complementary studies that investigated the impact of imputation on network reconstruction performance [80, 81]. In contrast to our study, we note that both these studies used experimental scRNA-seq data sets where it is usually difficult to determine the ground truth network. In [80], scRNA-seq data of seven different cell types were included, imputation methods such as MAGIC and SAVER as well as inference methods such as PIDC and GENIE3 were evaluated in this study. The authors found that MAGIC introduced high positive correlations and combining SAVER with PIDC led to an increase in network reconstruction performance, these findings are consistent with our results (see Supplemental Figures 2, 3, 4 and 5). However, some disagreements with our results were also observed. For example, while in this study it was reported that combining SAVER with PIDC gave better results than combining SAVER with GENIE3, we found these combinations of imputation method and network inference algorithm are comparable regardless of the network sparsity and topology (Supplemental Figures 3, 4 and 5). We also found that combining SAVER and GENIE3 does not improve the network inference precision over downsampled data (Supplemental Figures 3(C),(G),(K), 4(C),(G) and 5(C),(G)), unlike what was reported in the aforementioned study where the authors observed that combining SAVER and GENIE3 does improve network inference performance in some cases. In [81], it was reported that low capture efficiencies pose a challenge for imputation and network inference methods and that some imputation methods, namely DCA [23], preserve the gene-gene correlations structure even though false positive correlations are introduced, these findings are consistent with our results (Supplemental Figures 3, 4 and 5) where we found that for low capture efficiencies, regardless of the imputation and network inference method, the network inference precision is poor and we also found that SAVER similar to DCA preserves the gene-gene correlations structure as mentioned above.

In summary, biomodelling.jl uses mechanistic models of gene regulatory network and stochastic agent-based models of gene expression in cell populations to simulate realistic scRNA-seq data. This kind of approach is complementary to methods that are purely statistical and use deep neural networks (see e.g. [82]). As illustrated in this study, this kind of approach that is based on a known ground truth is useful for bench-marking and development of novel methods for the analysis of scRNA-seq data and gene-regulatory network inference.

## 5 Availability of data and materials

The raw datasets supporting the conclusions of this article are available in the following github repository: https://github.com/Msturroc/biomodelling_benchmark.

## 6 Authors contributions statement

A.L., V.S. and M.S. wrote the main manuscript text. A.L. prepared figures and performed simulations. V.S. and M.S. reviewed the manuscript.

## 7 Ethics and consent to participate

Not applicable to this study.

## 8 Consent to publish

Not applicable to this study.

## 9 Competing interests

The authors have no competing interests as defined by BMC, or other interests that might be perceived to influence the results and/or discussion reported in this paper.

## Acknowledgements

This work was supported by the European Union’s Horizon 2020 research and innovation programme under the Marie Skłodowska-Curie ITN initiative (Grant number: 766069).

## Supplemental material

### 1 Choosing threshold parameter for network in ference algorithms

**Supplemental Figure 1:**
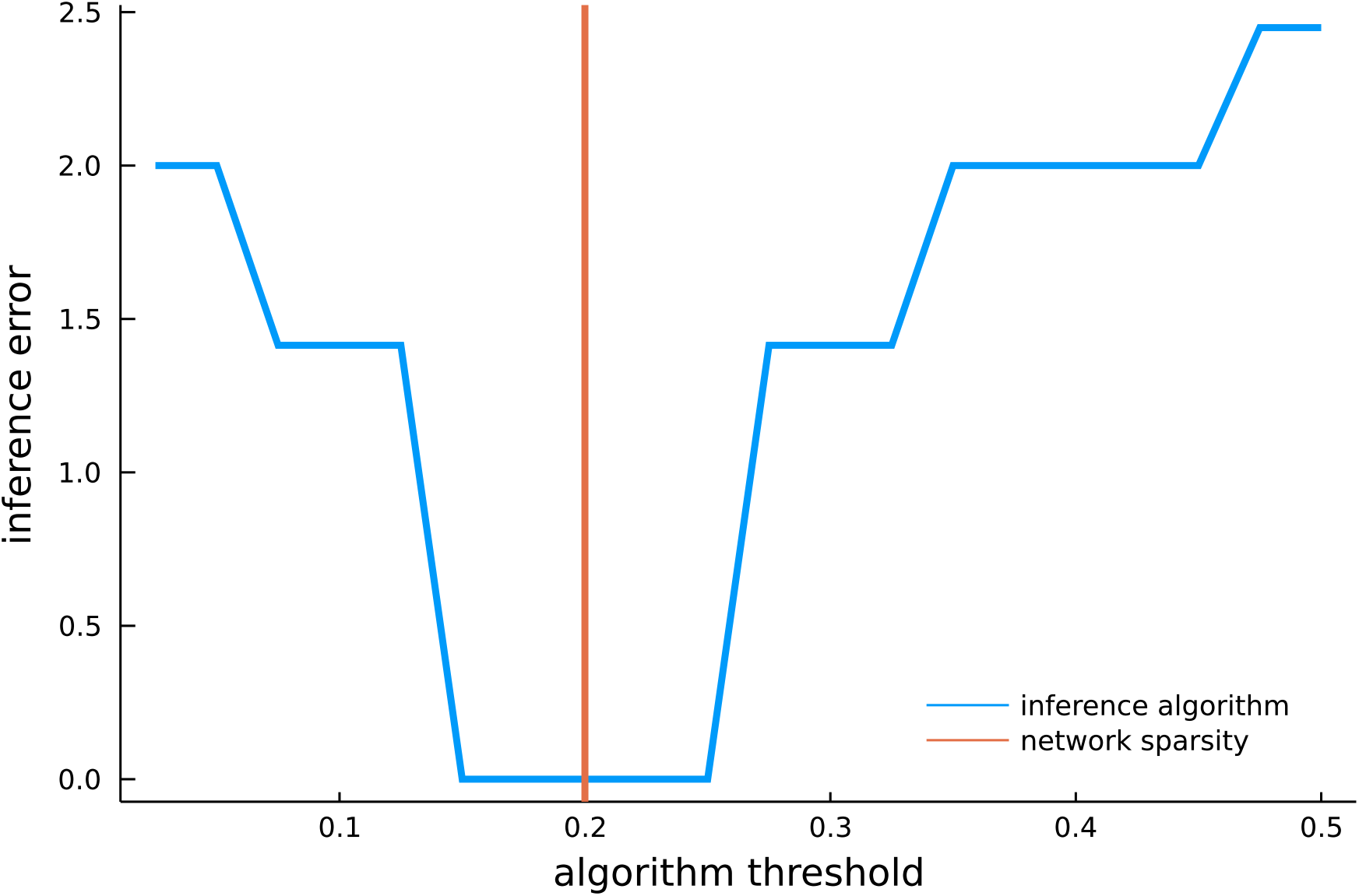
Impact of varying threshold parameter for network inference algorithms: plot showing PIDC inference error (as defined by the *l*_2_ norm of predicted adjacency matrix with ground truth adjacency matrix) as a function of the algorithm threshold for a 5-gene network toy example. The orange solid line is the network sparsity and the blue solid line represents the network inference algorithm error.

### 2 Example gene-gene correlations

**Supplemental Figure 2:**
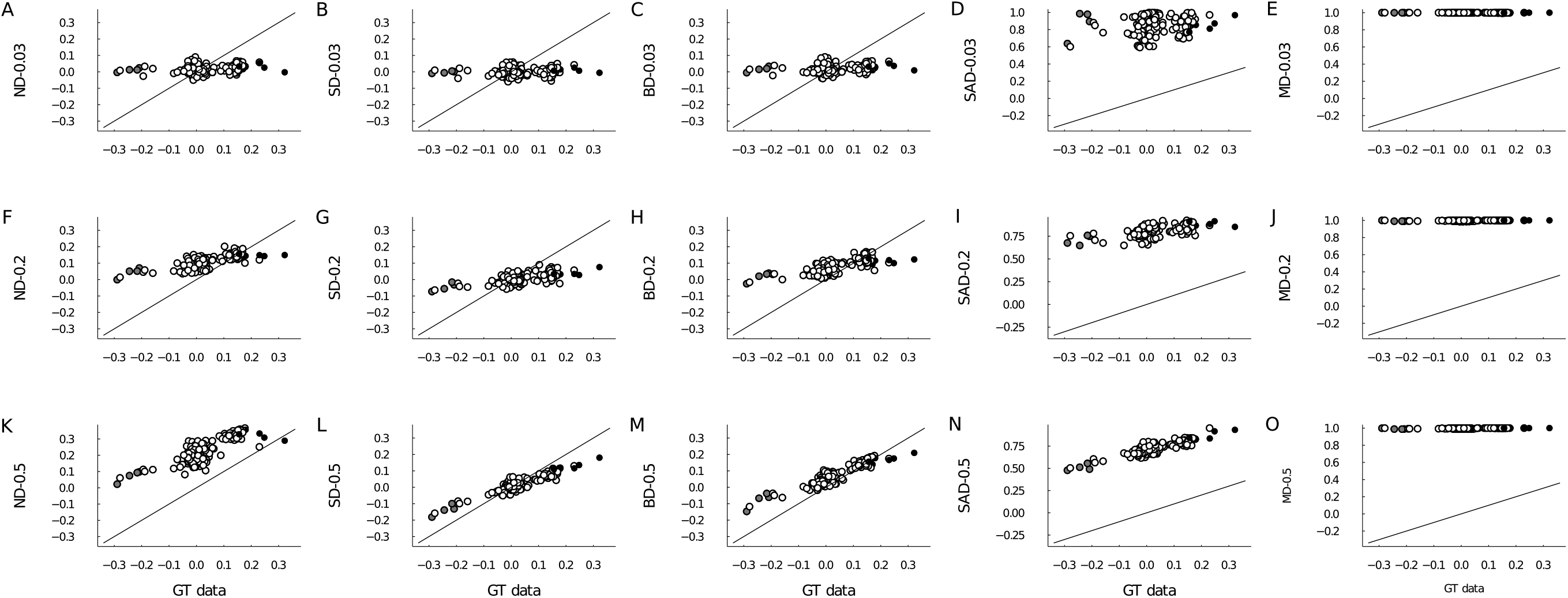
Impact of imputation on Pearson gene-gene correlations: results for capture efficiency rates 0.03, 0.2 and 0.5 are shown in the first, second and third row respectively and the imputation/scaling methods appear in the y-axis label of each plot. The x-axis show the gene-gene correlations in GT data. The black solid line represents the linear equation *x* = *y*, black dots are activation type reactions, grey dots are inhibition type reactions and open circles represent non-reacting type.

### 3 Inference precision: 20 gene additive regulation

**Supplemental Figure 3:**
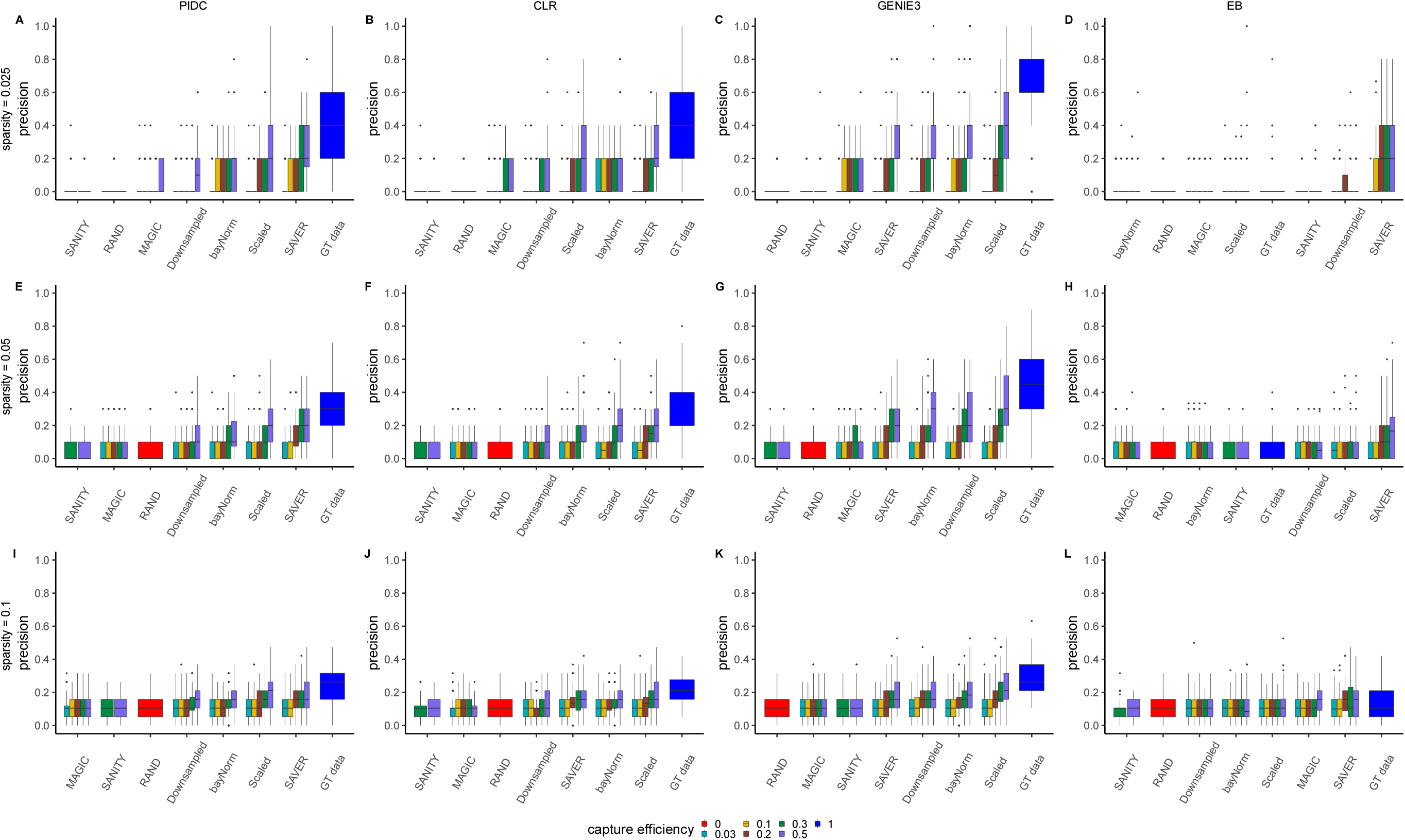
Impact of imputation on network inference performance for 100 different simulated 20 gene networks using sparsity = 0.025, 0.05 and 0.1 with **additive regulation** for various captures efficiency. RAND corresponds to precision obtained using random classification and GT data corresponds to precision obtained without downsampling (i.e., capture efficiency is set to 1). Plots (A) to (D) show box-plots of precision scores obtained for different imputation algorithms displayed on x-axes for PIDC, CLR, GENIE3 and Empirical Bayes respectively with sparsity = 0.025. Plots (E) to (H) show box-plots of precision scores obtained for different imputation algorithms displayed on x-axes for PIDC, CLR, GENIE3 and Empirical Bayes respectively with sparsity = 0.05. Plots (I) to (L) show box-plots of precision scores obtained for different imputation algorithms displayed on x-axes for PIDC, CLR, GENIE3 and Empirical Bayes respectively with sparsity = 0.1.

### 4 Inference precision: 20 gene multiplicative regulation

**Supplemental Figure 4:**
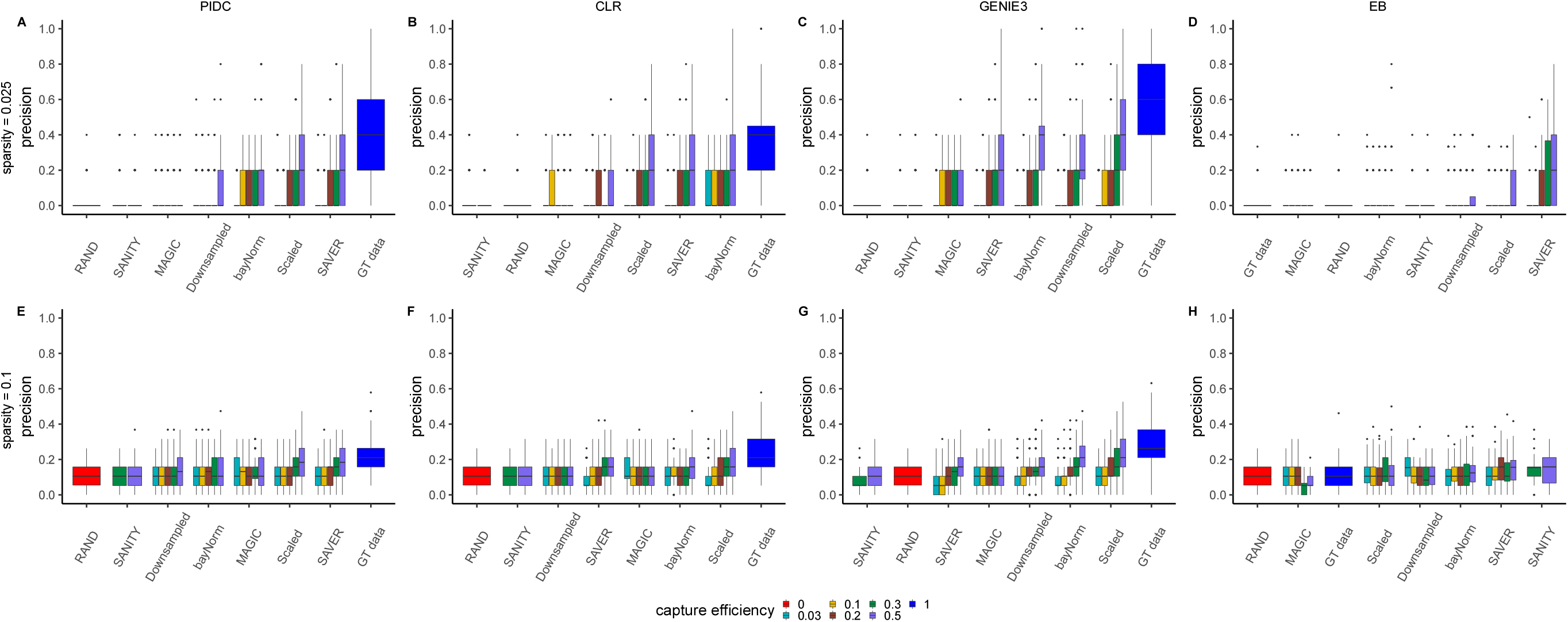
Impact of imputation on network inference performance for 100 different simulated 20 gene networks using sparsity = 0.025 and 0.1 with **multiplicative regulation** for various captures efficiency. RAND corresponds to precision obtained using random classification and GT data corresponds to precision obtained without downsampling (i.e., capture efficiency is set to 1). Plots (A) to (D) show box-plots of precision scores obtained for different imputation algorithms displayed on x-axes for PIDC, CLR, GENIE3 and Empirical Bayes respectively with sparsity = 0.025. Plots (E) to (H) show box-plots of precision scores obtained for different imputation algorithms displayed on x-axes for PIDC, CLR, GENIE3 and Empirical Bayes respectively with sparsity = 0.1.

### 5 Inference precision: 50 gene case, two different sparsity levels

**Supplemental Figure 5:**
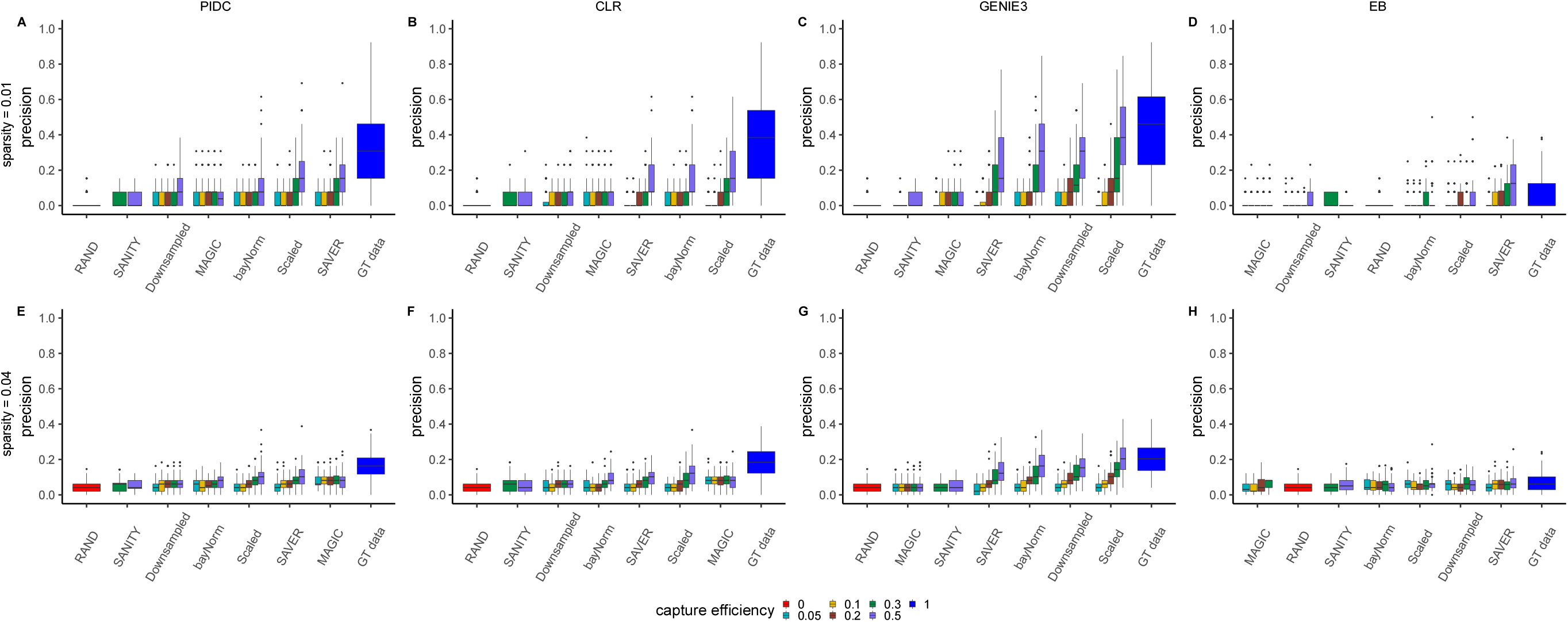
Impact of imputation on network inference performance for 100 different simulated 50 gene networks using sparsity = 0.01 and 0.04 with **multiplicative regulation** for various captures efficiency. RAND corresponds to precision obtained using random classification and GT data corresponds to precision obtained without downsampling (i.e., capture efficiency is set to 1). Plots (A) to (D) show box-plots of precision scores obtained for different imputation algorithms displayed on x-axes for PIDC, CLR, GENIE3 and Empirical Bayes respectively with sparsity = 0.01. Plots (E) to (H) show box-plots of precision scores obtained for different imputation algorithms displayed on x-axes for PIDC, CLR, GENIE3 and Empirical Bayes respectively with sparsity = 0.04.

### 6 Number of combination reactions in both 20 and 50 gene networks

**Supplemental Figure 6:**
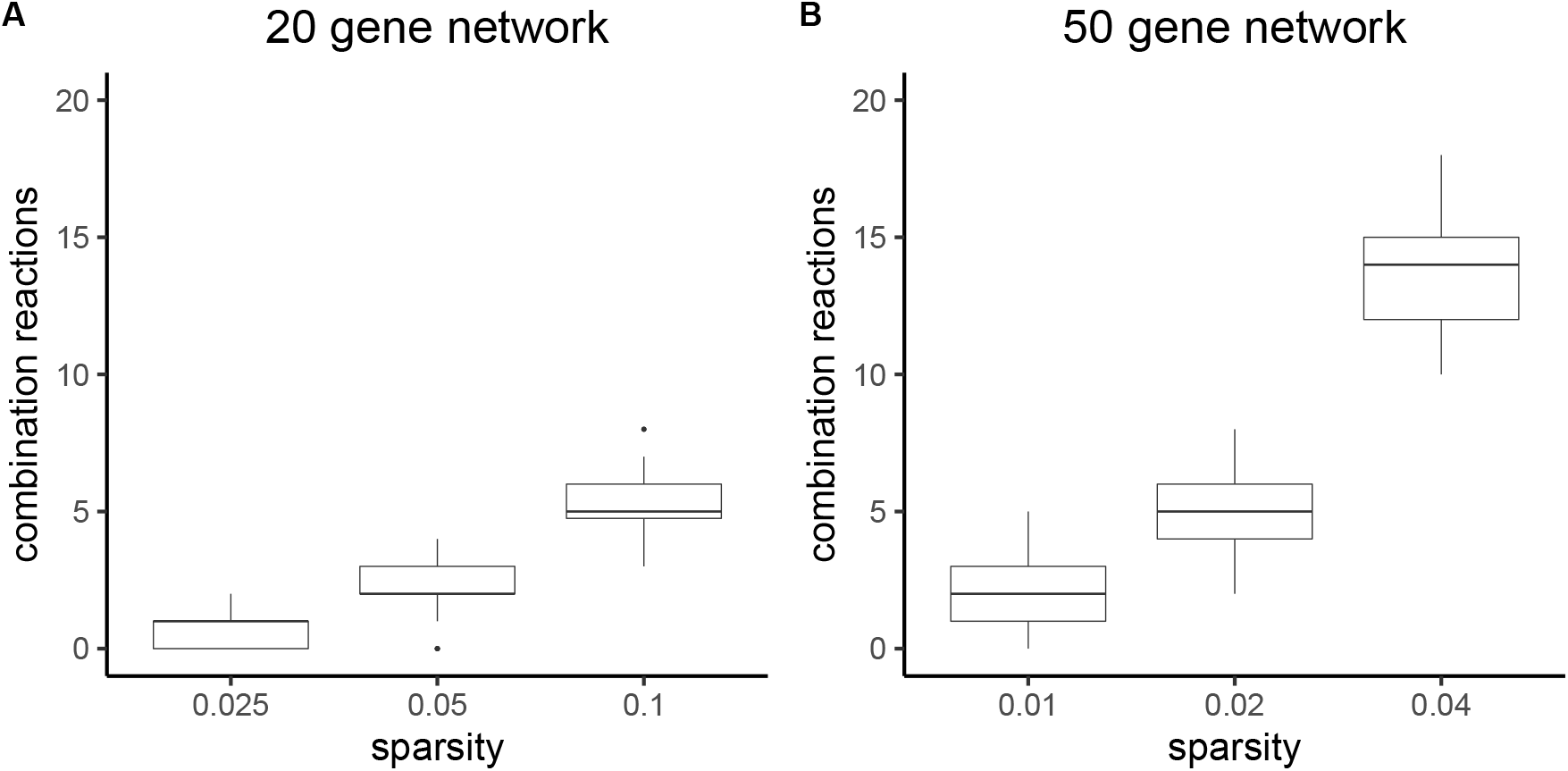
Number of combination reactions in 100 models of 20-genes and 50-genes networks for different sparsity levels. (A) box-plots of the number of combination reactions in 20-genes network for sparsity = 0.025, 0.05 and 0.1. (B) box-plots of the number of combination reactions in 50-genes network for sparsity = 0.01, 0.02 and 0.04.

### 7 Comparison of 20 gene network with 0.1 spar-sity and 50 gene network with 0.02 sparsity

**Supplemental Figure 7:**
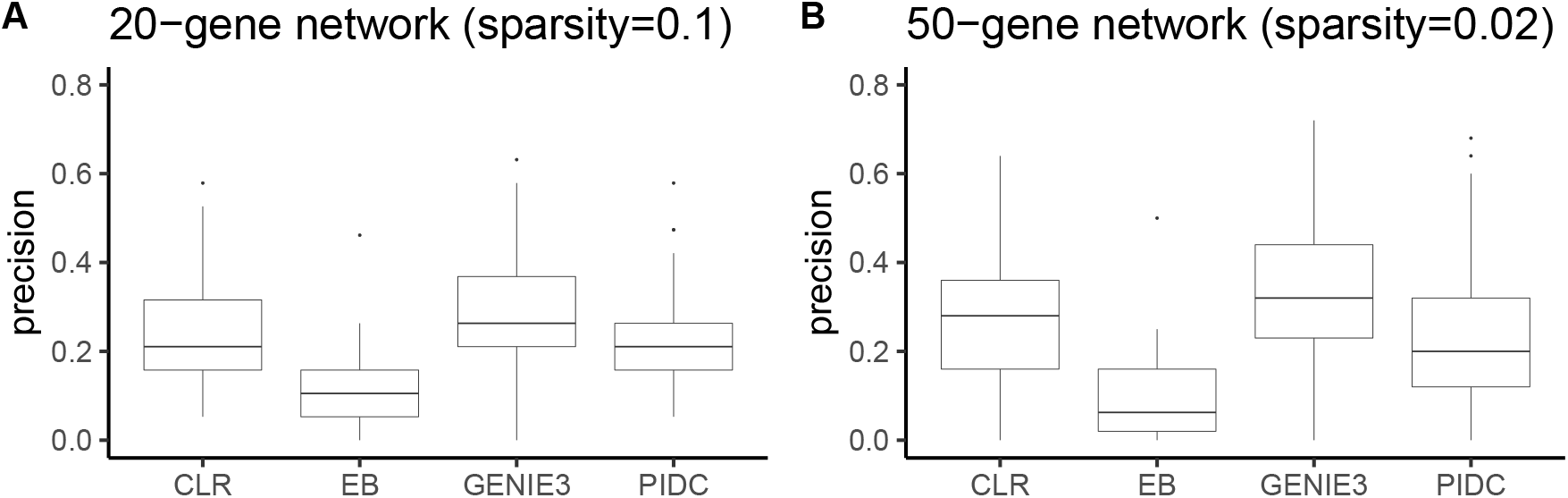
Performance of 100 models of 20-genes and 50-genes networks for 0.1 and 0.02 sparsity levels respectively. (A) box-plots of the precision for 100 20-genes networks with sparsity = 0.1. (B) box-plots of the precision for 100 50-genes networks with sparsity = 0.02.

## Footnotes

1* https://github.com/ayoublasri/Biomodelling.jl

2* https://github.com/ayoublasri/Biomodelling.jl/tree/master/parameters

## Notes

### Competing Interest Statement

The authors have declared no competing interest.

